# Restriction Enzyme Based Enriched L1Hs sequencing (REBELseq)

**DOI:** 10.1101/710095

**Authors:** Benjamin C. Reiner, Glenn A. Doyle, Andrew E. Weller, Rachel N. Levinson, Esin Namoglu, Alicia Pigeon, Gabriella Arauco-Shapiro, Emilie Dávila Perea, Cyndi Shannon Weickert, Gustavo Turecki, Deborah C. Mash, Richard C. Crist, Wade H. Berrettini

## Abstract

Long interspersed element-1 retrotransposons (LINE-1 or L1) are ~6 kb mobile DNA elements implicated in the origins of many Mendelian and complex diseases. The actively retrotransposing L1s are mostly limited to the L1 human specific Ta subfamily. In this manuscript, we present REBELseq as a method for the construction of differentially amplified next-generation sequencing libraries and bioinformatic identification of Ta subfamily long interspersed element-1 human specific elements. REBELseq was performed on DNA isolated from NeuN+ neuronal nuclei from postmortem brain samples of 177 individuals and empirically-driven bioinformatic and experimental cutoffs were established. REBELseq reliably identified both known and novel Ta subfamily L1 insertions distributed throughout the genome. Differences in the proportion of individuals possessing a given reference or non-reference retrotransposon insertion were identified. We conclude that REBELseq is an unbiased, whole genome approach to the amplification and detection of Ta subfamily L1 retrotransposons.

## Introduction

Retrotransposons are a class of mobile DNA elements capable of replicating and inserting copies of themselves elsewhere in the genome using an RNA intermediate (1). Long interspersed element-1 (L1), a type of retrotransposon that is currently active in the human genome, is estimated to constitute approximately 17% of the human genome (2). Despite their abundance, most L1s are truncated or mutated to the point of no longer being able to retrotranspose (3). There are ~100 L1s in each human genome that are competent, meaning they are full length and capable of replicating, the vast majority of which belong to the L1 human specific (L1Hs) Ta subfamily (4). Alternatively, in agreement with Kazazian *et al*. (5), Beck *et al.* described two competent L1s of the pre-Ta subfamily (2). A competent L1Hs is ~6 kb in length and contains a promoter, 5’ and 3’ untranslated regions, and two open reading frames (ORF): ORF1, encoding an RNA binding protein (6), and ORF2, encoding a fusion protein that acts as both a reverse transcriptase (7) and endonuclease (8).

Whereas competent retrotransposons are mostly limited to full length L1Hs of the Ta subfamily, the remnants of vast numbers of retrotransposons, from both evolutionarily older and Ta subfamily L1Hs can be found throughout the human genome. Notably, Ta subfamily L1Hs contain identifying nucleotides in the 3’ UTR that can be utilized for their differential amplification by polymerase chain reaction (PCR) (4). Retrotransposition of competent L1Hs elements frequently results in 5’ truncation of the new L1Hs element insertions. Therefore, to reliably amplify and detect full length and truncated L1Hs elements, including germline polymorphic and individual somatic mutations, from the Ta subfamily, PCR primers for amplification must be targeted near the 3’ end of the L1Hs Ta subfamily sequence.

In this manuscript, we present REBELseq (<u>R</u>estriction <u>E</u>nzyme <u>B</u>ased <u>E</u>nriched <u>L</u>1Hs <u>seq</u>uencing), a scalable technique for the differential amplification of both full length and 5’ truncated L1s. As described below, we specifically target the Ta subfamily of L1Hs and summarize the results of the application of REBELseq to DNA samples from 177 individuals.

## Materials and Methods

### Brain samples and purification of genomic DNA from NeuN+ nuclei

177 fresh frozen human postmortem prefrontal cortex brain samples (Brodmann’s Area 9 or 10) were provided by the Douglas-Bell Canada Brain Bank at McGill University, the Human Brain and Spinal Fluid Resource Center at UCLA, the Sydney Brain Bank at Neuroscience Research Australia, or the University of Miami Brain Endowment Bank. All work was approved by the University of Pennsylvania Institutional Review Board as category four exempt human subject research. All reported age, sex and ethnicity data are based on associated medical records. Sample set contained 138 males (Age 45.6 ± 15.5; 93 European, 25 African, 15 Hispanic, 2 Asian and 3 of unknown ethnic origin) and 39 females (Age 55.5 ± 19.2; 33 European, 1 African, 3 Hispanic and 2 of unknown ethnic origin). See Supplemental Methods for full details.

Brain samples were homogenized and co-stained with DAPI and an α-NeuN-AF488 antibody (Millipore # MAB377X) using a modification of a previously described method (9), and NeuN+ and NeuN-nuclei were isolated using a FACSAria II (Beckman-Coulter). Isolated nuclei were lysed overnight in digestion buffer in a 56 °C water bath, and the following morning genomic DNA (gDNA) was isolated using the Zymo Research genomic DNA clean and concentrator kit (#D4011). The concentration of gDNA was quantified using a Qubit 3 fluorometer and high-sensitivity double-stranded DNA assay (ThermoFisher #Q32854). See Supplemental Methods for full details.

### Ta subfamily L1Hs-enriched library construction and sequencing

Ta subfamily L1Hs-enriched next generation sequencing libraries were constructed utilizing the REBELseq technique (Fig. 1). 33 ng of gDNA extracted from NeuN+ neuronal nuclei was digested with HaeIII, in the presence of shrimp alkaline phosphatase. A single primer extension utilizing a 3’ diagnostic ‘A’ nucleotide (L1HsACA primer, see Supplemental Oligomers for all primer sequences; all oligos from IDT, Iowa, USA), a more stringent variation of a primer originally designed by Ewing and Kazazian (10), extends only Ta subfamily L1 3’ UTR sequence (4) and the adjacent downstream genomic DNA, leaving a terminal 3’ A-overhang. The A-overhang products were ligated to a double stranded T-linker molecule, originally designed for a different technique (11), and the ligated products were amplified using the L1HsACA primer and a T-linker specific primer to enrich the number of copies of each unique Ta subfamily L1Hs insertion (Primary PCR). The Primary PCR product was diluted and used as a template for a hemi-nested Secondary PCR reaction using the T-linker primer and the Seq2-L1HsG primer. The purpose of the hemi-nested secondary PCR reactions is three-fold: to reduce the length of 3’ L1 sequence carried forward, to add an Illumina sequencing primer, and, most importantly, to use the L1Hs diagnostic 3’ G nucleotide of the Seq2-L1HsG primer to further enrich the L1Hs Ta subfamily. Secondary PCR products were cleaned and size selected using KAPA Pure Beads (Roche #KK8002) according to the manufacturer’s protocol for a 0.55X-0.75X double-sided size selection, generating a purified library of 200-1,000 bp fragments. A Tertiary PCR using 5’ overhang primers then added Illumina flow cell adapters, containing a single multiplexing index, to both ends of the amplicon. Sequencing libraries then underwent a final cleanup and size selection using KAPA Pure Beads 0.55X-0.75X double-sided size selection and the average fragment size of the library was determined using the 2100 Bioanalyzer and high sensitivity DNA kit (Agilent #5067-4626). The quantity of sequenceable library for each sample was determined using the KAPA Library Quantification Kit (Roche #KK4835) on the 7900HT real-time PCR (Applied Biosystems). Barcoded samples were pooled in groups of 6, and pooled libraries were sequenced by the University of Pennsylvania Next-Generation Sequencing Core, using one pool per lane, on an Illumina HiSeq 4000 utilizing 150 bp paired-end sequencing. See Supplemental Methods for full details.

**Figure 1:**
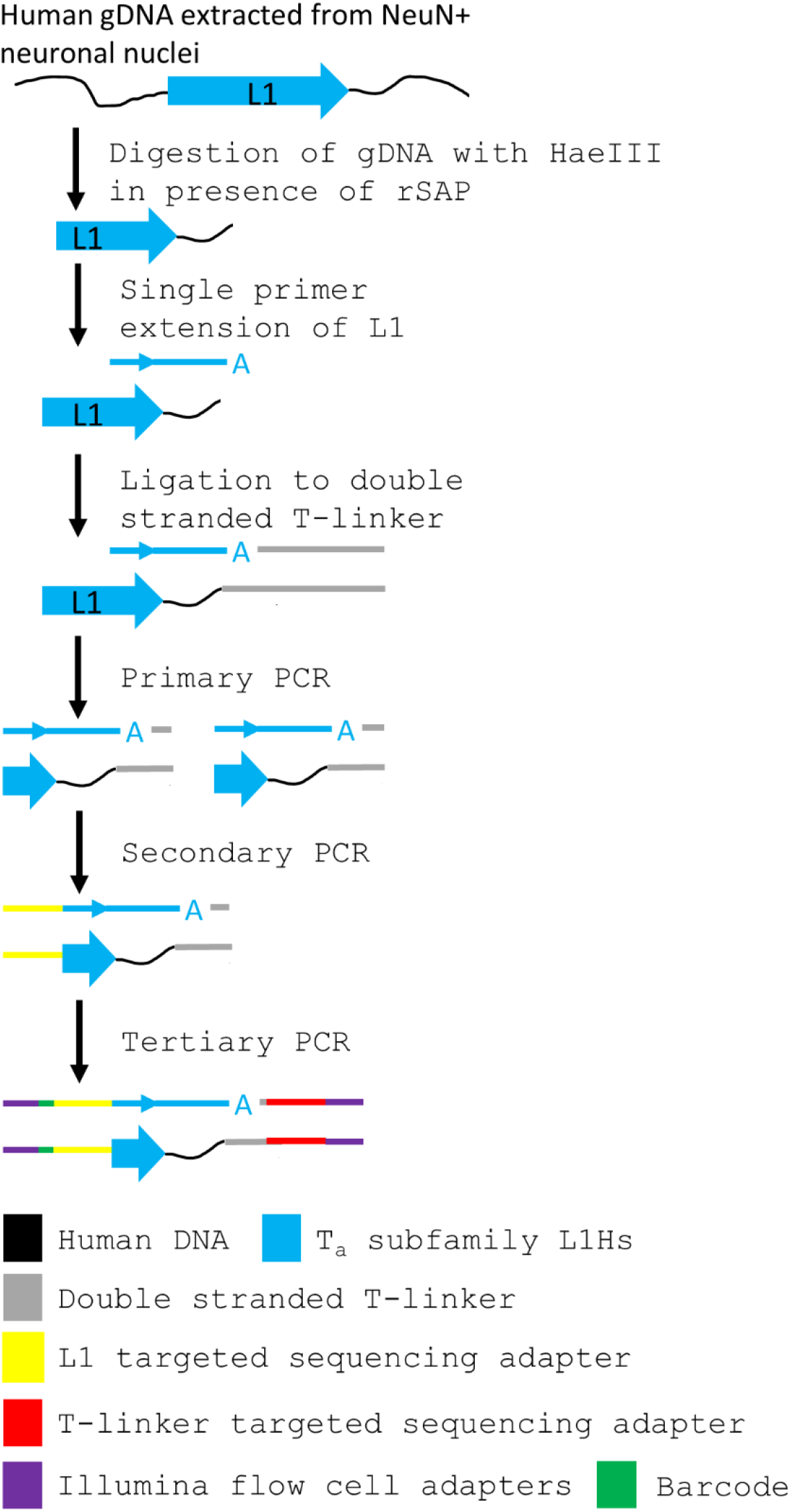
Schematic of the construction of Ta subfamily enriched L1Hs sequencing libraries. gDNA isolated from NeuN+ nuclei was enzymatically digested with HaeIII to fragment the genome. A single primer extension using Ta subfamily specific L1HsACA primer extends the 3’ end of the L1 sequence into the downstream gDNA. The 3’ ‘A’ overhang from the single primer extension is ligated to a custom T-linker, and primary PCR amplifies the construct using L1HsACA and T-linker specific primers. Hemi-nested secondary PCR using the L1Hs specific L1HsG primer and T-linker primer reduces the length of the L1 sequence carried forward and adds a sequencing adapter to the L1 end. Tertiary PCR uses 5’ overhang primers to add barcodes to the L1 end and Illumina flow cell adapters to both ends of the amplicon.

### Bioinformatics

Demultiplexed sequencing data was cleaned and quality trimmed to a Phred quality score of Q ≥ 20 using BBTools bbduk (https://sourceforge.net/projects/bbmap), and trimmed read 1 data was aligned to the hg19 build of the human genome with Bowtie2-2.1.0 (12) using end-to-end, very-sensitive alignment to generate a sequence alignment map (SAM) file for each individual. ~10% of unaligned reads contained a poly(T) sequence after a genomic sequence which made it unalignable. Assuming these poly(T) stretches likely corresponded to the 3’ poly(A) tail of the L1Hs sequence, they were trimmed, and the trimmed reads were aligned to hg19 with Bowtie2-2.1.0 using end-to-end, very-sensitive alignment parameters to generate a second SAM alignment file for each individual. SAM files were converted to BAM format, with reads having alignment MapQ scores < 30 being discarded, using samtools-0.1.19 (13). Using samtools-0.1.19, the two BAM files for each individual were combined. The resulting single BAM files from all individuals were then merged using RG tags, a tag that allows for the combination of reads from multiple BAM files, while still retaining the source (i.e. the individual) from which each aligned read was generated, and the merged file was sorted and indexed. The merged BAM file was stripped into BAM files for individual chromosomes using samtools-0.1.19, and the chromosome specific BAM files were used as input for the custom python script REBELseq_v1.0. REBELseq_v1.0 utilizes the sorted and indexed reads in the chromosome specific BAM files to generate peaks of overlapping reads within a 150 base pair sliding window, and does so with respect to DNA strand. Each peak corresponds to a putative L1 retrotransposon insertion, for which the peak’s genomic coordinates, number of unique reads per peak, number of reads per individual sample and average read alignment quality (mean MapQ) are determined. The REBELseq_v1.0 output file was then further annotated using the custom python script REBELannotate_v1.0 for L1Hs annotated in hg19 repeat masker and L1Hs identified in the 1000 genomes data (14). REBELannotate_v1.0 utilizes a browser extensible data (BED) formatted file to annotate genomic features of interest that overlap with or occur within 500 base pairs downstream, with respect to strand, of the peak being annotated. The REBELseq_v1.0 and REBELannotate_v1.0 custom scripts are based on work originally described by Ewing and Kazazian (10), and all python scripts and necessary reference files discussed in this manuscript are available online (https://github.com/BenReiner/REBELseq). The raw sequencing data that was analyzed to support the findings of this manuscript are available from the corresponding author upon request. See Supplemental Methods for full details.

### PCR Validations

Primers used in PCR experiments for method validation were designed using Primer3-2.2.3 in a Perl script (makeprimers.pl), originally written by Adam Ewing (10), with an optimal Tm setting of 58°C (minimum 56°C and max 63°C), an optimal primer length of 24 nucleotides (minimum 21 and max 27) and the GC-clamp option set to 1. 25 µL reactions were constructed using 1x Go-Taq colorless hotstart master mix (#M5133, Promega), 1 ng of gDNA for both an individual predicted to have a particular L1Hs insertion or an individual predicted not to have the insertion, and either 0.2 µM of the filled site and empty site (empty allele detection) or the filled site and L1HsACA primer (filled allele detection). Samples were thermally cycled as 95 °C 2:30 min, then 10 cycles of touchdown PCR at 95 °C 0:30 min, 69-60 °C 0:30 min, 72 °C 1:30 min, then 25 cycles of 95 °C 0:30 min, 60 °C 0:30 min, 72 °C 1:30 min, then 72 °C 5:00 min, 4 °C hold. Go-Taq green flexi buffer (#M8911, Promega) was added to all samples after amplification, and samples were separated by 1.2% gel electrophoresis and visualized on a GelDoc XR+ (Bio-Rad).

### Statistics

Data are presented as mean ± standard deviation.

## Results

### REBELseq identifies L1Hs retrotransposon insertions

We performed REBELseq using DNA isolated from postmortem cortex NeuN+ nuclei samples of 177 individuals and identified a total of 157,178 independent putative Ta subfamily L1Hs insertions, with 1,050 annotating to hg19 known reference L1Hs elements (ref L1Hs) and 156,128 not annotating to hg19 known reference L1Hs (non-ref L1Hs). The ref L1Hs we detected represented ~68% of all L1Hs annotated in hg19 repeat masker (1,050/1,544; see Discussion) and were distributed across all chromosomes (Fig. 2a). Of the non-ref L1Hs, 432 annotated to known L1s in the 1000 genomes data (known non-ref L1Hs), while the other 155,696 are putative novel non-ref L1Hs. The ref L1Hs had an average mean MapQ of 38.82 ± 4.10. Having high confidence that the detected ref L1Hs were real, we used the ref L1Hs average mean MapQ minus two standard deviations (MapQ ≥ 30.62) as a bioinformatic cutoff of our data, after which we retained 974 ref L1Hs (92.76% of the 1,050 ref L1s), 430 known non-ref L1Hs (99.54% of 432) and 127,976 putative novel non-ref L1Hs (82.20% of 155,696).

**Figure 2:**
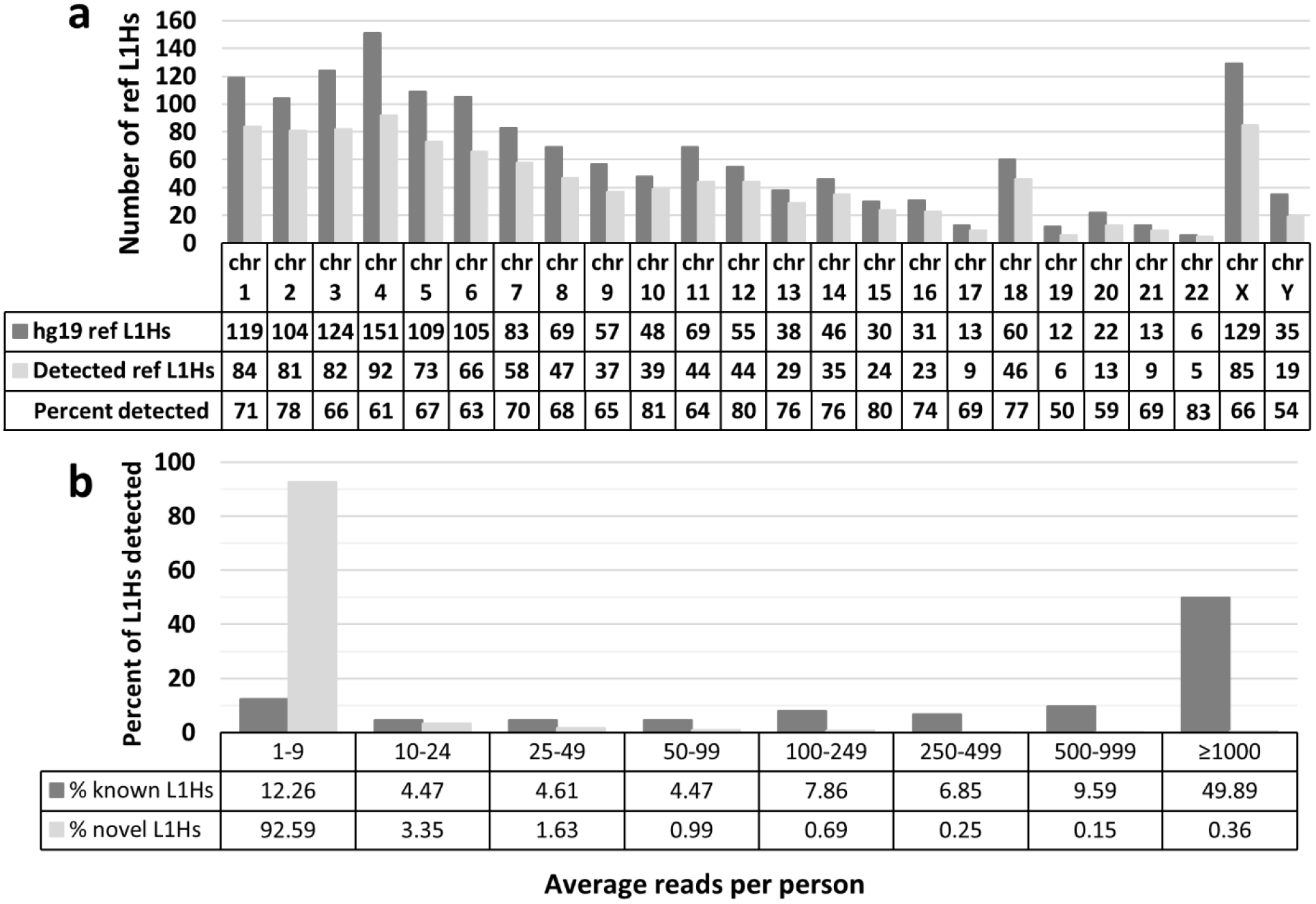
Detection levels of Ta subfamily L1Hs. **a** The number of ref L1Hs detected per chromosome versus the number of L1Hs annotated in hg19 repeat masker. Overall, we detected ~68% of L1Hs annotated in hg19 repeat masker, which aligns with our expectations based on the literature (see discussion). **b** Average number of sequencing reads per person for a given L1 insertion. Data was binned to show contrast between the average number of sequencing reads per person seen for known and putative novel L1Hs insertions. An average number of sequencing reads per person ≥ 100 was used as a bioinformatic cutoff.

In the remaining known (ref L1Hs and known non-ref L1Hs) and putative novel L1Hs, we next examined the average number of sequencing reads per person for a given L1Hs insertion (Fig. 2b), and a clear difference between the known and novel L1Hs emerged, with 92.6% of putative novel L1Hs having average reads < 10, and 87.7% of known L1Hs having average reads ≥ 10 and 74.2% having average reads ≥ 100. Having confidence in the known L1Hs pool, an average of ≥ 100 sequencing reads per person was used as a bioinformatic cutoff, which retained 699 ref L1Hs (66.57%), 332 known non-ref L1Hs (76.85%) and 1,842 putative novel non-ref L1Hs (1.18%).

### L1Hs insertion validation by PCR

Having established bioinformatic cutoffs for our data, we next sought to experimentally determine the proportion of bioinformatically-detected putative L1Hs insertions that were real using PCR of the 3’ insertion junction of the allele containing the insertion (filled allele) and the corresponding genomic region on the ‘empty’ allele (see Fig. 3 for example image). Having confidence that the known L1Hs insertions (ref L1Hs and known non-ref L1Hs) are real because they were previously reported, we focused on the validation of the putative novel non-ref L1Hs insertions predicted by REBELseq. The percentages of putative novel non-ref L1Hs insertions that could be validated by average read number bin were experimentally determined, and these data were used to calculate the number of detected putative novel non-ref L1Hs insertions that we would expect to be true positives in each read bin. Using the number of putative novel non-ref L1Hs insertions predicted to be true positives and the numbers of known L1Hs detected, we calculated the probability of validating any detected L1Hs insertion per read bin and the cumulative probability of validating any detected L1Hs insertions using the lower value of a read bin as a minimum cutoff for the data (Table 1).

**Figure 3:**
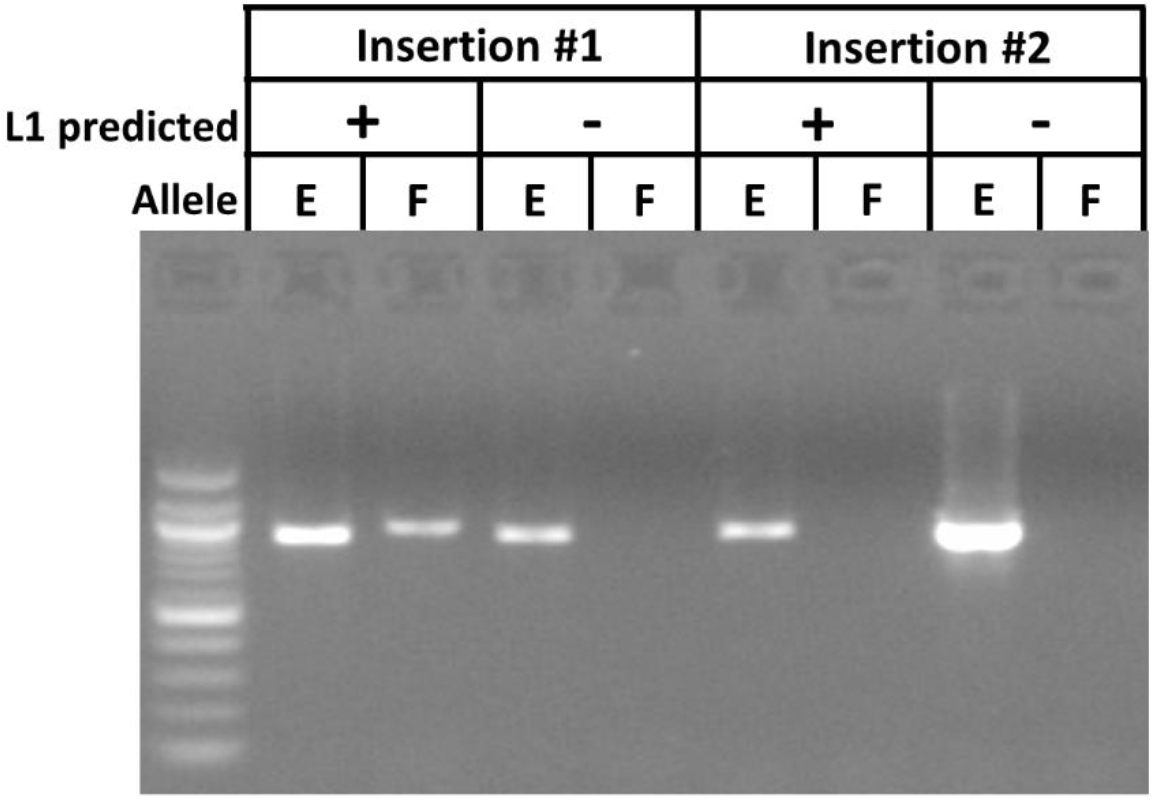
Example image of a successful and unsuccessful confirmatory PCR. PCR experiments were conducted to determine the proportion of putative novel non-ref L1Hs insertions detected by REBELseq that could be independently validated. For each insertion, a person predicted to have the L1Hs insertion (+) and a person not predicted to have the insertion (-) were used to amplify the genomic region purported to contain the L1Hs insertion (Filled site, F) and the same genomic region if it did not contain the insertion (Empty site, E). Insertion #1 is a positive confirmation of the results predicted by REBELseq, while Insertion #2 is a negative confirmation.

**Table I:** Table describing the rates of putative L1Hs insertion validation by average number of reads. The percentage of successfully validated L1Hs per read bin was used to calculate the number of real novel non-ref L1Hs. The percent validated per bin was calculated as the sum of real novel non-ref L1Hs, number of detected known non-ref L1Hs and number of detected ref L1Hs, divided by the sum of the number of detected novel non-ref L1Hs, number of detected known non-ref L1Hs and number of detected ref L1Hs. The cumulative probability of being validated was calculated similarly, we the bottom of the lowest included read bin serving as a lower cutoff.

| Avg. #<br>of reads | novel non-ref L1Hs |  |  |  |  | known<br>non-ref<br>L1Hs | ref L1Hs | %<br>validated<br>per read<br>bin | cumulative<br>probability<br>of being<br>validated<br>(%) |
| --- | --- | --- | --- | --- | --- | --- | --- | --- | --- |
|  | #<br>Successful | #<br>Attempted | %<br>Successful | #<br>detected | #<br>real | # detected | #<br>detected |  |  |
| ≥1,000 | 32 | 44 | 72.7 | 460 | 334 | 227 | 465 | 89.1 | 89.1 |
| 500-999 | 16 | 41 | 39.0 | 186 | 73 | 43 | 90 | 64.6 | 83.8 |
| 250-499 | 13 | 40 | 32.5 | 320 | 104 | 28 | 67 | 48.0 | 75.9 |
| 100-249 | 6 | 36 | 16.7 | 876 | 146 | 34 | 77 | 26.0 | 58.8 |

### Distribution of detected L1Hs

The genomic distribution of known and novel L1Hs, per 10 MB genomic window, of our highest quality L1Hs detection data was ascertained (≥ 1,000 average reads, Fig. 4a), and the distribution of the number of individuals having a given known (Fig. 4b) or novel (Fig. 4c) L1Hs for these data was assessed.

**Figure 4:**
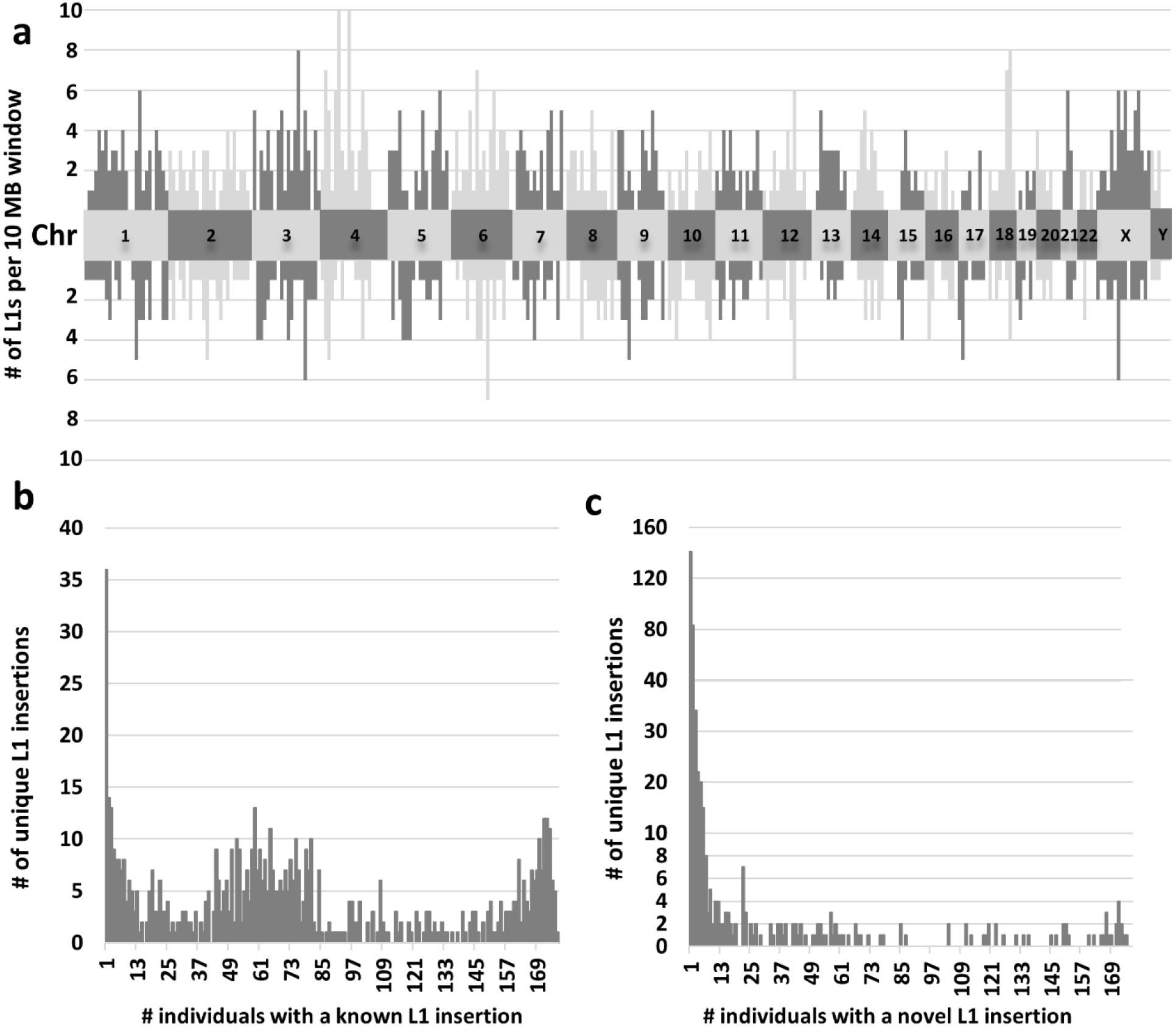
Distributions of high confidence L1Hs insertions. **a** The genomic distribution of known L1Hs (above the chromosome numbers) and novel L1Hs (below the chromosome numbers) per 10 MB window of each chromosome. Alternating color pattern and labeled central blocks represent the different chromosomes. Known and novel L1Hs are distributed throughout the genome, suggesting REBELseq is an unbiased whole genome approach. **b** Number of individuals sharing a known L1Hs insertion. The number of known L1Hs insertions shared by different numbers of individuals shows a trimodal distribution. While some ref L1Hs occur with a rate of affected individuals approaching 1.0, the other local maxima are focused at few or less than half of surveyed individuals. This demonstrates that known L1Hs should be considered polymorphic in nature, rather than ubiquitous, in the human genome. **c** Number of individuals sharing a novel L1Hs insertion. The number of novel L1Hs insertions shared by different numbers of individuals shows a right skewed distribution. While some novel L1Hs were detected in most individuals, most novel L1Hs were detected in one or a few individuals.

## Discussion

While other methods for preparation of L1-targeted next generation sequencing libraries exist (10, 11, 15–22), some are labor-intensive when scaled to a large number of individual genomes, because of reliance on gel purification (10, 11) or because they require preparation of multiple amplicon libraries per individual sample, due to the use of multiple hemi-degenerate primers (10, 19). Some methods rely on the random shearing of DNA (18, 22, 23), a process that requires specialized equipment and somewhat abundant DNA quantity, while others may have reduced specificity for the Ta subfamily of L1Hs, which constitutes the majority of actively replicating L1 retrotransposons in the human genome (16, 20, 21, 23). REBELseq was designed as a high throughput alternative method to reliably detect the 3’ flanking region of Ta subfamily L1Hs elements with limited gDNA input. It should be noted that REBELseq was not intended to determine whether an L1Hs insertion is full length or truncated or if ORF1 and ORF2 are intact, but merely to determine the presence or absence of the Ta subfamily L1Hs. After identification of insertions of interest, these other aspects can be determined by long range PCR and Sanger sequencing.

Other methods to produce L1-enriched libraries employing restriction enzyme digests of gDNA as a starting point have been described (15). There is no set requirement for which restriction enzyme to use in REBELseq other than that it generates blunt ends and cuts 5’ of the L1HsACA primer that identifies the 3’ end of the Ta subfamily L1Hs elements. REBELseq begins with restriction enzyme digestion of gDNA with HaeIII. This restriction enzyme was chosen for the following reasons: 1) it creates blunt ends at 5’-GG|CC-3’ sequences such that polishing of cleaved ends is unnecessary; 2) it cuts the human genome to an average size of 342 ± 478 base pairs (NEBcutter) which is similar to the range of fragments generated by sonication; 3) it does not cut within the 3’ end of the L1Hs sequence targeted by our L1Hs Ta subfamily-specific primers. One possible concern of using a single restriction enzyme digestion in REBELseq library construction is the possibility that the restriction site for the enzyme occurs immediately downstream from a L1 insertion, thus preventing the single primer extension into the downstream gDNA (see Fig. 1). While this raises the possibility that using a single restriction enzyme digestion during REBELseq library construction may not detect all L1 insertions, it does not reduce the validity of L1 insertion that are detected. One possible solution to this concern would be to perform restriction enzyme digestions with two or more enzymes that meet the above criteria and pool the fragments before the single primer extension step.

Our method is specifically designed to leverage diagnostic nucleotides specific to the Ta subfamily of L1Hs elements for differential amplification. It can easily be adapted to detect the 5’ flanking region of full length L1Hs elements in the human genome, although diagnostic nucleotides are limited within the 5’ end of L1Hs for the Ta subfamily. Reliable primers targeting the 5’ end of Ta subfamily L1Hs and an enzyme other than HaeIII would be required for analysis of full length L1Hs elements, as HaeIII makes a single digestion in the L1Hs sequence near the 5’ end. REBELseq could also be utilized to differentially amplify and identify evolutionarily older pre-Ta subfamily L1Hs insertions (by changing the last three nucleotides of the L1HsACA primer to ‘ACG’) or any type of L1Hs element (by eliminating the last nucleotide of the L1HsACA primer to end in ‘AC’). Additionally, this method could be utilized to amplify and identify almost any other genomic element with a specific pair of nested primers (to replace L1HsACA and L1HsG).

The REBELseq technique is compatible with human gDNA purified from any tissue source, including gDNA purified from leukocytes in saliva and blood. In the application of REBELseq presented in this manuscript, 177 fresh frozen human postmortem prefrontal cortex brain samples were utilized for purification of gDNA from NeuN+ neuronal nuclei. Utilizing gDNA from NeuN+ neuronal nuclei allows for the identification of germline L1s, similar to using gDNA from a peripheral source, and individual neuronally relevant somatic mutations occurring: only in the brain (an L1 that retrotransposed early in neuronal development), in a limited number of neurons (an L1 that retrotransposed in a neuronal progenitor cell, producing a cluster of neurons harboring the insertion), or in a single neuron (an L1 that retrotransposed in the neuron post-mitotically), results which could be verified by comparing the REBELseq results from NeuN+ nuclei to those of a peripheral tissue. Using REBELseq to compare Ta subfamily L1Hs from multiple brain regions would also allow for the identification of regionally specific L1Hs insertions, possibly relevant in the context of neurological and psychiatric disease.

Using REBELseq, we identified 157,178 independent putative Ta subfamily L1Hs insertions, with 1,050 annotating to reference L1Hs in hg19 repeat masker. The ref L1Hs detected represent ~68% of all hg19 repeat masker L1Hs (Fig. 2a). The L1Hs elements annotated in hg19 repeat masker represent both Ta subfamily L1Hs, which were specifically targeted, and pre-Ta subfamily L1Hs. Thus, we did not expect to detect all L1Hs annotated in hg19 repeat masker. Previous work using the genomes of 15 individuals aligned to hg18 showed that the pre-Ta subfamily of L1Hs represented ~36% of L1Hs insertions, with Ta subfamily L1Hs elements composing the remainder (~64%) which closely approximates the number we detected (10).

The bioinformatic cutoffs described in this manuscript were empirically derived by the average mean MapQ score for the ref L1Hs, the average number of sequencing reads per individual per insertion (Fig. 2b), and experimental confirmation of putative novel non-ref L1Hs insertions (Table 1). We were unable to experimentally validate all putative novel L1Hs in any of our average read bins. This is possibly due to the detected L1Hs insertion being sequencing reads of a chimeric PCR product, or possibly because Ta subfamily L1Hs frequently retrotranspose into repetitive sequences in the genome (e.g. the remnants of older repetitive elements), making both their genomic alignment and PCR validation difficult. We believe this suggests the proportion we were able to validate likely represents a minimum value, with additional detected insertions being confirmable with more complex PCR methods. ~70% of all known L1Hs averaged at least 100 sequencing reads per person, with our experimentally determined probability of validating any detected L1Hs above 100 reads being ~59%. Examining the data with ≥1,000 reads per person, we still see ~47% of all detected known L1Hs and a probability of validating any detected L1Hs of almost 90%. These data suggest that novel non-ref L1Hs having ≥1,000 reads per person include what are likely real polymorphic and somatic L1Hs insertions.

When examining the genomic distribution of known and novel L1Hs having an average of ≥1,000 sequencing reads per person (Fig. 4a), we see that both the known and novel L1Hs are dispersed throughout the genome, suggesting that REBELseq is an unbiased whole genome approach. Assessing the distribution of the number of individuals having either a known (Fig. 4b) or novel (Fig. 4c) L1Hs insertion, we observe that both groups have L1 insertions occurring throughout the range of possible values (i.e. number of individuals). With respect to the known L1Hs, there appears to be a trimodal distribution with ~37% of L1Hs occurring in 45-85 individuals and ~9% occurring in ≤ 3 and ≥ 170, meaning ~55% of insertions occur in only ~29% of the possible numbers of individuals. Our finding that a large proportion of known L1Hs insertions occur in a small fraction of individuals disagrees with a previous report (10), but we believe this may be due to their small sample size, thereby diminishing the likelihood that an individual would not have a given insertion. ~70% of novel L1Hs occur in ≤ 7 people, while ~8% of novel L1Hs occur in more than two thirds of individuals, suggesting that these L1Hs insertions likely represent uncatalogued ref L1Hs occurring at common minor allele frequencies.

Taken together, we believe these data demonstrate that REBELseq reliably detects both known and novel Ta subfamily L1Hs insertions, and that this technique could be a powerful method for examining the association of the prevalence of L1Hs insertions in a broad range of human disease.

## Acknowledgements

The authors wish to thank those providing support for this work. BR is supported, in part, by a 2017 NARSAD Young Investigator Grant (#26634) from the Brain and Behavior Research Foundation as the Patrick A. Coffer Investigator, funding for which was generously provided by Ronald and Kathy Chandonais. BR was supported during part of this work by T32 MH014654 (PI is WB). This work was funded by National Institute of Health grants to WB (R21NS095756, R01DA040972 & R01MH109260). RC was supported by K01DA036751. Additionally, the authors wish to thank all the participants, and their families, who donated their brain tissue to research for making these studies possible.

