## Supplemental Data for "Restriction Enzyme Based Enriched L1Hs sequencing (REBELseq)"

### Supplemental Methods

Brain Samples

177 fresh frozen human postmortem prefrontal cortex brain samples (Brodmann's Area 9 or 10) were provided by the Douglas-Bell Canada Brain Bank at McGill University, the Human Brain and Spinal Fluid Resource Center at UCLA, the Sydney Brain Bank at Neuroscience Research Australia, or the University of Miami Brain Endowment Bank. All samples were from sudden death cases without prolonged agonal conditions. Causes of death involving neural trauma or asphyxiation and post-mortem intervals > 48 hours were excluded.

Purification of DNA from NeuN+ nuclei

Brain samples were processed and stained for NeuN by modification of a previously described method (1). Frozen brain samples (40-80 mg) were homogenized on wet ice using a Dounce homogenizer (Sigma #D8938-1SET) in 1 mL sucrose lysis buffer, containing 0.32 M sucrose, 5 mM $CaCl_2$, 3 mM $MgOAc_2$, 0.1 mM EDTA, 10 mM Tris, 1 mM DTT, 1 mM PMSF, 15 µL protease inhibitor cocktail (Sigma #P8340), 5 µL RNaseOUT (Invitrogen #10777019) and 1 µL Trition X-100, using 25 strokes with the large clearance pestle and 25 strokes with the small clearance pestle. 500 µL aliquots of homogenate were layered on top of 200 µL of sucrose cushion solution, containing 1.8 M sucrose, 3 mM $MgOAc_2$, 10 mM Tris, 1 mM DTT, 1 mM PMSF, 2 µL protease inhibitor cocktail and 2.5 µL RNaseOUT, in a 1.5 mL Eppendorf style tube and centrifuged at 21 130 x g for 30 minutes (min) at 4 °C to pellet nuclei. The middle protein layer and upper liquid phase, containing total cytosolic RNA, were removed, diluted in 4 mL of TRIzol solution (Invitrogen #15596-018) and frozen at -80 °C for future analysis. The remaining sucrose cushion solution was removed and discarded, taking care not to disturb the nuclei pellet. 65 µL of PBS + 3 mM $MgCl_2$ was added to each nuclei pellet and incubated on wet ice for 15 min, to allow the pellet to soften. Following incubation, pellets were further loosened by gentle up and down pipetting. The solutions containing each pair of pellets, from the previously separated homogenate, were transferred to a 2 mL Eppendorf style tube for nuclei staining, with

55 µL PBS + 3 mM $MgCl_2$, 25 µL BSA blocking solution, containing 0.5% BSA and 10% normal goat serum, 10 µL 7.5 µM DAPI and 30 µL 1:10 diluted α-NeuN-AF488 antibody (Millipore #MAB377X). Nuclei staining reactions were incubated on a nutator, protected from light, at 4 °C for 60 min. Staining solutions were loaded onto Falcon tubes with 30 µm cell strainer caps (Fisher #08-771-23) and filtered by gravity flow.

Labeled nuclei were sorted into NeuN+ and NeuN- fractions in 15 mL conical vials containing 500 µL of PBS + 3 mM $MgCl_2$ cushion using a FACSAria II (Beckman-Coulter; See Supplemental Methods Figure 1 for example FACS separation image). After sorting, NeuN+ and NeuN- nuclei fractions were diluted to 2 mL final volume using PBS + 3 mM $MgCl_2$. 400 µL (0.2 volumes) of sucrose cushion solution, described above, was added to each tube, mixed by gentle inversions and allowed to incubate, on ice, for 15 min. Diluted nuclei fractions were centrifuged at 2,000 x g for 30 min at 4 °C to pellet nuclei, after which, all but 100 µL of supernatant was removed by careful pipetting. 600 µL digestion buffer, containing 50 mM Tris, 100 mM EDTA, 100 mM NaCl and 1% SDS, and 35 µL proteinase K solution (Sigma #3115828001) was added, mixed by vortexing and allowed to incubate, overnight, in a 56 °C water bath. The following morning, NeuN+ and NeuN- genomic DNA (gDNA) were isolated using the Zymo Research genomic DNA clean and concentrator kit (#D4011), utilizing the manufacturer's protocol, and the concentration was determined using a Qubit 3 fluorometer and high-sensitivity double-stranded DNA assay (ThermoFisher #Q32854).

Construction of Ta subfamily enriched L1Hs amplified sequencing libraries

33 ng (~5000 diploid genomes) of NeuN+ gDNA for each sample was digested with 5 U HaeIII (New England Biolabs #R0108M) in a 10 µL reaction, containing 1X CutSmart buffer (New England Biolabs #B7204S) and 1 U recombinant shrimp alkaline phosphatase (New England Biolabs #M0371S) according to the manufacturer's protocol  (HaeIII digestion, Fig. 1). Samples were incubated at 37 °C 60:00 min, then 80 °C 20:00 min, then 4 °C hold. All the

digested NeuN+ gDNA was then used as a template for a 25 µL single cycle primer extension reaction (Single Primer Extension, Fig. 1), containing 1X GoTaq HotStart Colorless Master Mix (Promega #M5133) and 33.2 fmol/µL L1HsACA primer (See Supplemental Oligomers), which should extend only Ta subfamily L1 3' UTR sequence (2) and the adjacent downstream genomic DNA. Samples were incubated at 95 °C 2:30 min, then 95 °C 0:30 min, 60 °C 1:00 min, 72 °C 1:00 min, then 4 °C hold. All of the L1HsACA primed, 3' A-overhang extension products were ligated to a double stranded T-linker molecule (See Supplemental Oligomers), with a 5' phosphate on the top strand, in a 30 µL reaction (T-linker ligation, Fig. 1), containing 1X T4 DNA Ligase Buffer (New England Biolabs #B0202S), 400 U T4 DNA Ligase (New England Biolabs #M0202L) and 2.78 pmol/µL double stranded T-linker. Samples were incubated at 16 °C for 120 min, then 65 °C 20 min, then 4 °C hold. All of the T-linker ligated, Ta subfamily L1 elements were amplified (Primary PCR, Fig. 1) to enrich the number of copies of each unique Ta subfamily L1 in a 50 µL reaction, containing 1X GoTaq HotStart Colorless Master Mix, 0.2 pmol/µL L1HsACA primer and 0.2 pmol/µL T-linker bottom strand primer (See Supplemental Oligomers). Primary PCR was thermal cycled as 95 °C 2:30 min, then 20 cycles of 95 °C 0:30 min, 60 °C 0:30 min, 72 °C 1:30 min, then 72 °C 5:00 min, 4 °C hold. The primary PCR product was then diluted 1:10 in nuclease free water and used as template in a 25 µL hemi-nested PCR reaction (Secondary PCR, Fig. 1), containing 2.5 µL 1:10 diluted Primary PCR, 1X GoTaq HotStart Colorless Master Mix, 0.2 pmol/µL T-linker bottom strand primer and 0.2 pmol/µL Seq2-L1HsG primer (See Supplemental Oligomers). Secondary PCR was thermally cycled as 95 °C 2:30 min, then 35 cycles of 95 °C 0:30 min, 60 °C 0:30 min, 72 °C 1:30 min, then 72 °C 5:00 min, 4 °C hold. The secondary PCR product was cleaned and size selected using KAPA Pure Beads (Roche #KK8002) according to the manufacturer's protocol for a 0.55X-0.75X double-sided size selection, generating a 12 µL eluate of purified library of 200-1,000 bp fragments. The purified secondary PCR library was then used as a template for the addition of one of six single indexed Illumina flow cell adapters in a 50 µL reaction (Tertiary

PCR, Fig. 1) , containing 11 µL purified secondary PCR library, 1X Phusion HF Polymerase Buffer (New England Biolabs #B0518S), 2.5 U Phusion Hot Start HF polymerase (New England Biolabs #M0535L), 0.2 mM dNTPs (Thermo #R0191), 0.5 *p*mol/µL Adapt2-Seq1 primer and 0.5 *p*mol/µL Adapt1-Barcode-Seq2 primer (See Supplemental Oligomers). Tertiary PCR was thermally cycled as 98 °C 0:30 min, then 15 cycles of 98 °C 0:05 min, 72 °C 0:15 min, then 72 °C 1:00 min, 4 °C hold. Finally, the tertiary PCR reactions underwent final cleanup and size selection by KAPA Pure Beads 0.55X-0.75X double-sided size selection, with a final eluant volume of 22 µL. This purification was verified, and the average fragment size of the library was determined, using the 2100 Bioanalyzer and high sensitivity DNA kit (Agilent #5067-4626; see Supplemental Methods Figure 2 for example bioanalyzer trace).

Library quantification, pooling and sequencing

The barcoded, L1-enriched sequencing libraries were quantified, in triplicate, for the concentration of PCR fragments properly flanked by the Illumina flow cell adapters using the KAPA Library Quantification Kit (Roche #KK4835) and 7900HT real-time PCR (Applied Biosystems) according to the manufacturer’s protocol, utilizing 1:20,000, 1:200,000 and 1:2,000,000 dilutions. Using the KAPA Library Quantification Analysis Template_ILM_v4-1, the Ct values and the average fragment size (determined above by Bioanalyzer), the concentration of sequenceable library was calculated for each sample. Samples were pooled in groups of six, each having a unique barcode from the Adapt1-Barcode-Seq2 primer (described above), in 30 µL pools of 25 nM equimolar final concentration. Pooled libraries were then sequenced by the University of Pennsylvania Next-Generation Sequencing Core, using one pool per lane, on an Illumina HiSeq 4000 utilizing 150 bp paired-end sequencing and 10% PhiX sequencing control spike-in (Illumina #FC-110-3001). Raw sequencing data was evaluated by Illumina quality filters and reads passing filter were deconvoluted by sequencing lane and barcode and transferred into read 1 and read 2 (paired end) FASTQ files.

Bioinformatics

Demultiplexed FASTQ files from sequencing were transferred to the Penn Medicine Academic Computing Services servers for analysis. Initial evaluation of sequencing read quality showed high quality reads for read 1 and diminished quality for read 2 (paired end reads), likely due to the need to sequence through the low diversity L1HsG primer and the low complexity, highly repetitive, variable length poly(A) tail of L1Hs elements. Therefore, only read 1 sequence data was used for all subsequent bioinformatics. PhiX spike-in sequences were removed from an individual's FASTQ file using BBTools (https://sourceforge.net/projects/bbmap) bbduk (BBTools decontamination using k-mers) and the PhiX sequence as reference. Remaining reads were end trimmed of Illumina flow cell and sequencing adapters and cleaned based on a Phred quality score of $Q \geq 20$ using bbduk. Trimmed reads were aligned to the hg19 build of the human genome with Bowtie2-2.1.0 (3) using the end-to-end, very sensitive alignment parameters, to generate a sequence alignment map (SAM) file for each individual. Investigation into the unaligned reads showed that ~10% contained a poly(T) stretch after the genomic portion of the read that was unalignable, possibly corresponding to the poly(A) sequence at the 3' end of non-reference L1Hs elements. As such, bbduk was used to k-trim poly(T) stretches from the 3' end of the unaligned reads using a k-mer of 11 T's and a hamming distance of 1. The poly(T) trimmed reads were realigned to hg19 with Bowtie2-2.1.0 using end-to-end, very sensitive alignment parameters, generating a second SAM alignment file for each individual. Each SAM file was converted to a binary alignment map (BAM) format, during which alignments with a MapQ score < 30 were removed using samtools-0.1.19 (4). The two BAM files for each individual, one from each alignment, were merged, sorted and indexed using samtools-0.1.19. Using samtools-0.1.19, BAM files from all individuals were merged into a single BAM file using RG tags (-r option) and this combined BAM file was sorted and indexed. The single merged BAM file with RG tags was split into BAM files containing the data for each chromosome using samtools-0.1.19, and the chromosome specific BAM files were used as input for the custom python script REBELseq_v1.0. REBELseq_v1.0 utilizes the sorted and indexed reads in the chromosome specific BAM files to generate peaks of overlapping reads within a 150 base pair sliding window, and does so with respect to DNA strand. Each peak corresponds to a putative L1 retrotransposon insertion, for which the peak's genomic coordinates, number of unique reads per peak, number of reads per individual sample and average read alignment quality (mean MapQ) are determined. The REBELseq_v1.0 output file was then further annotated using the custom python script REBELannotate_v1.0 for L1Hs annotated in hg19 repeat masker and L1Hs identified in the 1000 genomes data. REBELannotate_v1.0 utilizes a browser extensible data (BED) formatted file to annotate genomic features of interest that overlap with or occur within 500 base pairs downstream, with respect to strand, of the peak being annotated. All REBELseq_v1.0 peaks that were annotated by REBELannotate_v1.0 as an L1Hs in hg19 repeat masker are referred to throughout the manuscript as reference L1Hs. All peaks that did not annotate to an hg19 repeat masker L1Hs are collectively referred to as non-reference L1Hs. Non-reference L1Hs that annotate to the 1000 genomes data are referred to as known non-reference L1Hs and non-reference L1Hs that did not annotate to the 1000 genomes data are putatively referred to as novel non-reference L1Hs. The novel non-reference L1Hs include what are likely real L1Hs insertions (because they meet our cutoff criteria) that inserted into older repetitive elements (Alu, L1Pa, etc.) within the reference genome. The REBELseq_v1.0 and REBELannotate_v1.0 custom scripts are based on work originally described by Ewing and Kazazian (5).

Scrutiny of the output data from REBELseq showed a low level of index hopping between multiplexed samples. Index hopping was remediated by removing samples with less than 6% (6) of the maximum read count per set of six multiplexed samples for a given L1 insertion. It is notable that index hopping could be eliminated by utilizing a second unique bar code in tertiary PCR (i.e. unique dual-indexing), thereby causing index hopped reads to be discarded during demultiplexing (7).

**Supplemental Methods Figures**

Supplemental Methods Figure 1

FACS of NeuN stained nuclei. Representative image showing FACS separation of NeuN+ and NeuN- DAPI stained nuclei. Boxes indicate gating for populations of nuclei that were collecting for gDNA isolation.

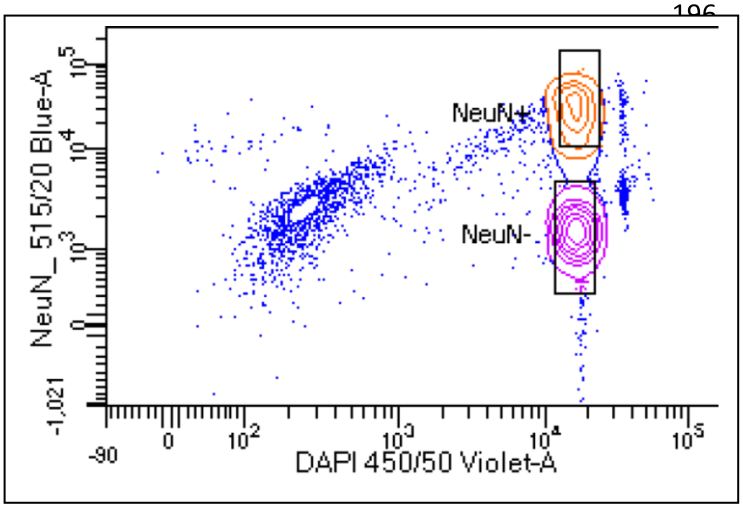

Supplemental Methods Figure 2

Confirmation of amplicon size selection. Representative bioanalyzer trace showing the size distribution of the size selected final, Ta subfamily-enriched L1Hs-amplified sequencing library for an individual sample. Size selected amplicon products were distributed between 200-1000 base pairs (bp, x-axis). Y-axis is arbitrary florescent units (FU).

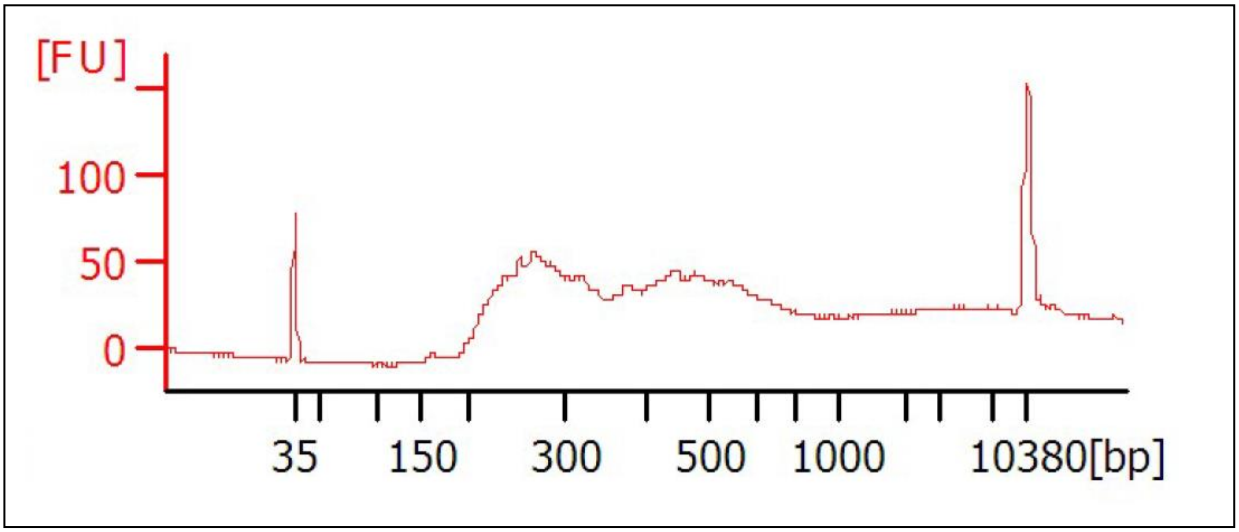

#### **Supplemental Oligomers**

L1HsACA primer
GGGAGATATACCTAATGCTAGATGACACA

Double stranded T-Linker
AGATGTGAGAAAGGGATGTGCTGCGAGAAGGCTAG/Phos/5’
TCTACACTCTTTCCCTACACGACGCTCTTCCGATC*T

T-linker bottom strand primer
TCTACACTCTTTCCCTACACGACGCTCTTCCGATCT

Seq2-L1HsG primer
GTGACTGGAGTTCAGACGTGTGCTCTTCCGATCTGCACATGTACCCTAAAACTTAG

Adapt2-Seq1
AATGATACGGCGACCACCGAGATCTACACTCTTTCCCTACACGACGCTCTTCCGATCT

Adapt1-Barcode-Seq2 - Index 1
CAAGCAGAAGACGGCATACGAGATACATCGGTGACTGGAGTTCAGACGTGTGCTCTTCCGATC*T

Adapt1-Barcode-Seq2 - Index 2
CAAGCAGAAGACGGCATACGAGATTGGTCAGTGACTGGAGTTCAGACGTGTGCTCTTCCGATC*T

Adapt1-Barcode-Seq2 - Index 3
CAAGCAGAAGACGGCATACGAGATCACTGTGTGACTGGAGTTCAGACGTGTGCTCTTCCGATC*T

Adapt1-Barcode-Seq2 - Index 4
CAAGCAGAAGACGGCATACGAGATATTGGCGTGACTGGAGTTCAGACGTGTGCTCTTCCGATC*T

Adapt1-Barcode-Seq2 - Index 5
CAAGCAGAAGACGGCATACGAGATGATCTGGTGACTGGAGTTCAGACGTGTGCTCTTCCGATC*T

Adapt1-Barcode-Seq2 - Index 6
CAAGCAGAAGACGGCATACGAGATTACAAGGTGACTGGAGTTCAGACGTGTGCTCTTCCGATC*T

* denotes a phosphorothioate bond and highlighting denotes barcode sequence
